# Exploration of Flavonoids to identify Potential Therapeutic Compounds by Targeting the Non-Structural V protein of Nipah Virus

**DOI:** 10.1101/2024.07.29.605559

**Authors:** S Rehan Ahmad, Md. Zeyaullah, Abdullah M. AlShahrani, Mohammad Suhail Khan, Haroon Ali, Khursheed Muzammil, Ali Mohieldin, Abdelrhman AG Altijani, Adam Dawria, Awad Osman Abdalla Mohamed, Abul Kalam

## Abstract

Researchers are interested in a number of interdisciplinary approaches that might speed up and reduce the cost of creating new medications. This work aims to determine target proteins and choose a lead medication to combat the Nipah Virus. Following a study of the literature, we discover the non-structural V protein (UniProt ID: Q997F2). I-TASSER was used to estimate the 3D structure. We examined the flavonoid dataset in search of a strong inhibitor. Pyrx is used to use AutoDock Vina for docking. The interactions between the drug and the target protein binding were examined using BIOVIA Discovery Studio. Desmond’s Molecular Dynamics Simulation (MD simulation) was used to study the stability of protein and inhibitor complexes in a physiological environment. Based on our research, we have designed two lead compounds that lessen the virus’s effect. This discovery will benefit science as it may lead to the development of novel medications. The newly discovered natural compounds showed promise as inhibitors, showing less adverse effects and more efficiency.

## Introduction

Nipah virus is a highly pathogenic and a bat-borne disease (generally transmitted from bats to humans) also characterized as a BSL-4 (Biosafety level 4) due to its high rate of virulence and lack of vaccines and therapies. It can also be transmitted through intake of contaminated food or directly from person-to-person. NiV is a negative-sense, single-stranded, non-segmented RNA and causes deadly encephalitis in human and also cause severe respiratory infections. Pteropus bats (flying foxes) are the natural reservoir of NiV. Nipah virus belongs to the genus Henipavirus, family Paramyxoviridae, order Mononegavirales. Fruit bats are natural hosts and they an asymptomatic not showing any symptoms of illness. First discovered during the outbreak in Singapore and Malaysia in 1999. Since then, it frequently causes outbreak in Southeast Asia, especially in Bangladesh. The case fatality rate of Nipah virus infection range from 40% to 75% [1-3].

Hendra virus (HeV) was the first henipavirus, discovered as zoonotic disease. This occurred in Australia back in 1994. In Australia there’s outbreak of severe illness in horses. Two people who were in close contact with horses also became ill, and due to illness one of them died. Scientist isolated a new paramyxovirus from both the human and horse’s cases, and done further experiments that confirmed that this virus was the cause of outbreak, later named as Hendra virus.

Hendra virus found in Australian flying foxes. When it spills over from bats to horses, it causes a severe respiratory and neurological illness [4, 5].

In March 1999, Nipah virus was isolated from the Sungai Nipah Village from the cerebrospinal fluid of patient. The outbreak causes 283 cases of symptoms and 109 deaths. 11 cases and one death were reported from Singapore among the workers of pig farms. Thes outbreaks are mostly due to close contact with pigs. In Malaysia large number of animals are raised together in slaughter houses, where the transmission of virus is more likely. Control strategies was successful in controlling the outbreaks like culling the pigs. However, there is no evidence of transmission of virus from human-to-human during these outbreaks [6, 7].

During September, December 1998 to January 1999, there were three types of outbreaks occur in Perak state of Malaysia near Ipoh city, town of sikamat of state Negri Sembilan, Malaysia and the largest outbreak in neighboring areas of Bukit Pelandok, Malaysia. All of these outbreaks were considered JEV a Japanese encephalitis virus which occur mostly in these areas. JEV related to yellow fever dengue and is spread by mosquitoes is a flavivirus (positive strand RNA virus, family Flaviviridae) [8, 9].

Nipah virus is the type of virus that belongs to family of viruses called Paramyxoviridae. It has a single strand of RNA and its genetic material is enclosed in lipid envelope. The negative sense RNA is arranged in helical symmetry. The RNA genome of Nipah virus is not divided into separate segments. Nipah virus encode six structural and three non-structural proteins. Structural protein includes Nucleoprotein (N), Phosphoprotein (P), Glycoprotein (G), fusion Glycoprotein (F), Matrix protein (M), Long polymerase (L). Within the Phosphorus gene, the alternative start codon results in the creation of small protein C. insertion of one or more no templated G residues result in the production of V and W protein by the editing of P mRNA [6, 10, 11].

Nipah virus takes 4 to 21 days to show symptoms after infection. It causes encephalitis which is severe swelling of brain and also breathing problems. Some people who get infected don’t show any symptoms at all. At first, people that were infected with NiV feels like they have flu, with a fever, headache, and myalgia (muscle pain). Within a week brain swelling signs appear. The most common symptoms include changes in mental state, weak muscles, jerky movements, difficulty in moving the eye, and weakness in limbs. Patient get worse quickly, go to coma and dying within days [12, 13].

If someone survive from Nipah virus, they might still have lasting problems with their brain and nerves, like feeling tired all the time or trouble in moving some part of body. Outbreaks causes some breathing problems, this breathing problem might show up as a cough, or lung problems. Nipah virus results in liver problems, brainstem issues, seizures, low platelet counts, being older and having other health problems.

Nipah virus (NiV) is a highly pathogenic disease with high fatality rate (40% to 70%) and causes severe diseases (respiratory and neurological). It indicates to critical public health issues due to its massive outbreaks all over the world, yet there are no FDA approved vaccines were available that can give an effective treatment to population against this virus. Traditionally methods to make a vaccine against Nipah virus takes a lot of resources and time consuming because they involve lot of trials and error to make effective vaccine. There is an urgent need to identify an effective treatment against NiV faster by using the computational methods (chemoinformatics, bioinformatics etc.) to reduce the illness rate and overcome the further outbreaks of Nipah virus [14-16].

This research aims to develop a computational method to design a best drug candidate against Nipah virus (NiV). This computational technique helps us to provide a best and faster lead candidate against Nipah virus (NiV) and be prepared for outbreaks [17].

## Materials and Methods

### Target retrieval from RCSB PDB and preparation

Using the targeted proteins’ unique UniProtKB ID: Q997F2 for the Nipah virus’s non-structural protein the sequence of the proteins was retrieved [18]. Proteins, DNA, and RNA are just a few of the many macromolecules for which the UniProtKB acts as a free online resource and data library [19, 20]. We used loop refinement using MODELLER to the protein structures. The proteins’ crystal structures were then optimized and minimized using RAMPAGE and Swiss PDB Viewer. To evaluate the quality of the protein structure, RAMPAGE in particular produced a Ramachandran Plot that showed no disagreements. Which amino acid residues were located in the preferred, permitted, or outlier zones was another piece of information this plot revealed. Additionally, we predicted the binding sites on the protein targets using CASTp (Computed Atlas of Surface Topography of Proteins) 3.0. CASTp is a reliable and comprehensive method for measuring and analyzing the topography of proteins, which helps to identify potential binding sites [21-24].

### Molecular Docking

From the literature, we chose a large number of the druggable flavonoids. The screening process employed AutoDock Vina, a molecular docking program [25, 26]. The chosen compounds were docked with certain protein receptors to assess their interactions and binding affinities with the target proteins. In order to do this, we used PyMOL to create complex files that included the receptor and ligand structures. Then, we used BIOVIA Discovery Studio to analyze and display the two-dimensional protein-ligand interactions. This helped us understand more about how these compounds interacted with the receptor proteins [27, 28].

### Lead Identification

We conducted an ADMET (Absorption, Distribution, Metabolism, Excretion, and Toxicity) analysis on the chosen medications using Qikprop. This technique simplified our assessment of the compounds’ diverse pharmacokinetic and pharmacodynamic properties to ascertain their suitability as potential therapeutic options. We discovered a particular chemical that, when combined with the docking binding affinity data and the results of the ADMET investigation, demonstrated intriguing qualities. To verify this molecule’s binding and stability with the target protein in a physiologically relevant setting, we conducted further biophysical and biochemical experiments on it. More knowledge was gained about the compound’s potential as a therapeutic option as well as its ability to successfully interact with the target protein in conditions that are similar to the biochemical environment of the human body.

### Molecular Dynamics Simulation

We performed 100 nanosecond molecular dynamics simulations using the Desmond program to get understanding of the dynamic behavior of the molecular system. The molecular dynamics (MD) simulations were initiated after completing an essential preliminary step known as first protein-ligand docking. Thanks to docking, the first static picture of the ligand’s binding inside the target protein’s active site was created. In this work, the movement and interactions of individual atoms during a certain time period were simulated using molecular dynamics models. In doing so, they provide a prediction of the ligand-binding behavior in a biologically relevant environment.

This dynamic modeling technique allows us to see the interactions and changes that occur between the molecules over time, providing us with a greater understanding of how the system functions in a fluid and dynamic environment [29-32].

We used Maestro’s Protein Preparation Wizard to optimize and fine-tune the ligand-receptor combination, which reduced the system and fixed any missing residues. The System Builder tool was used for the whole system building process. We used the OPLS_2005 force field and the TIP3P (Intermolecular Interaction Potential 3 Points Transferable) fluid model while keeping the temperature at 310 K and the pressure at 1 atm. We simulated a concentration of 0.15 M sodium chloride and added neutralizing ions to guarantee the system’s neutrality and mimic physiological circumstances. The models were first balanced and checkpoints were set up for frequent evaluation, with data being captured every 100 ps, before the modeling itself began [33, 34].

## Results and Discussion

The UniProtKB was the source of the receptor’s three-dimensional structure. Plays a critical function in inhibiting host immune responses. Blocks the generation and signaling route of interferon-alpha/beta, which prevents the creation of a cellular antiviral state. Interacts with host IFIH1/MDA5 and DHX58/LGP2 to block the transduction pathway involved in the activation of the IFN-beta promoter, shielding the virus from cell antiviral responses. Blocks the type I interferon signaling cascade by interacting with host STAT1 and STAT2, preventing their phosphorylation and subsequent nuclear translocation. Effectively inhibits the type II interferon signaling pathway. Interferon induction is suppressed by interacting with and stabilizing host UBXN1, which is a negative regulator of both RIG-I-like receptors (RLR) and the NF-kappa-B pathway [35, 36].

The picture shows the final protein structure after loop reduction and optimization, as well as the predicted binding site. The structure showed a strong degree of agreement with desired structural observations and an outstanding overall quality score of 90%. Said another way, Figure 1 is a graphic depiction of a protein’s three-dimensional structure predicted by I-TASSER, sequence taken from UniProtKB entry Q997F2. The region of the protein where it is anticipated to interact with other molecules is highlighted in this picture, which displays the structure of the protein following loop modifications and minimization. With 90% accuracy, the construction is of high quality and perfectly aligned with desired structural features. Researchers looking into the function and possible connections of the protein usually find this information to be significant.

**Figure 1:**
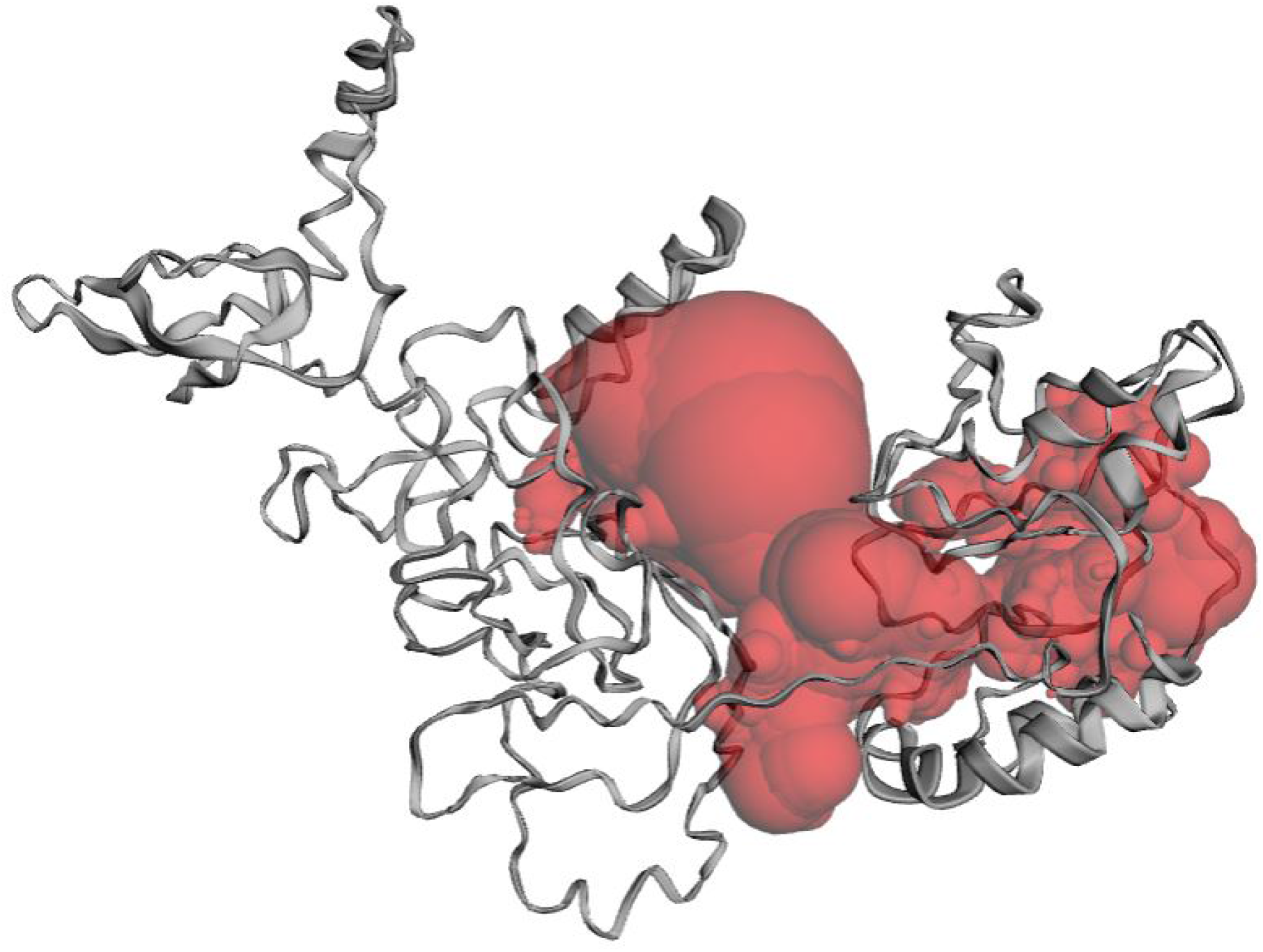
This figure illustrates the three-dimensional structure of a protein predicted by I-TASSER.

The top 5 leads’ binding affinities and ADMET analyses are listed in Table 1. All five compounds were shown to be the most active for the protein target after lead identification. The 2D interactions of these high-performing compounds are shown in Figure 2. Table 1 provides an overview of the best compounds’ characteristics. A molecular dynamics (MD) simulation lasting 100 nanoseconds was carried out for the protein targets in association with the most effective compound (CID_452707), in order to learn more about these intriguing chemicals. The MD trajectories were subjected to a number of studies, such as evaluations of protein-ligand interactions and root mean square deviation (RMSD and RMSF). For the top hits, docking investigations were also conducted using AutoDock Vina. In order to assess the pharmacokinetic and safety characteristics of these substances, an ADMET (absorption, distribution, metabolism, excretion, and toxicity) research was carried out utilizing instruments such as QikProp and pkCSM. As shown in Table 1, the top 5 compounds were chosen for additional review based on their ADMET and docking findings. The discovery of possible therapeutic candidates with both good pharmacophore matches and encouraging biological activity is made possible by this all-encompassing method.

**Table 1:**
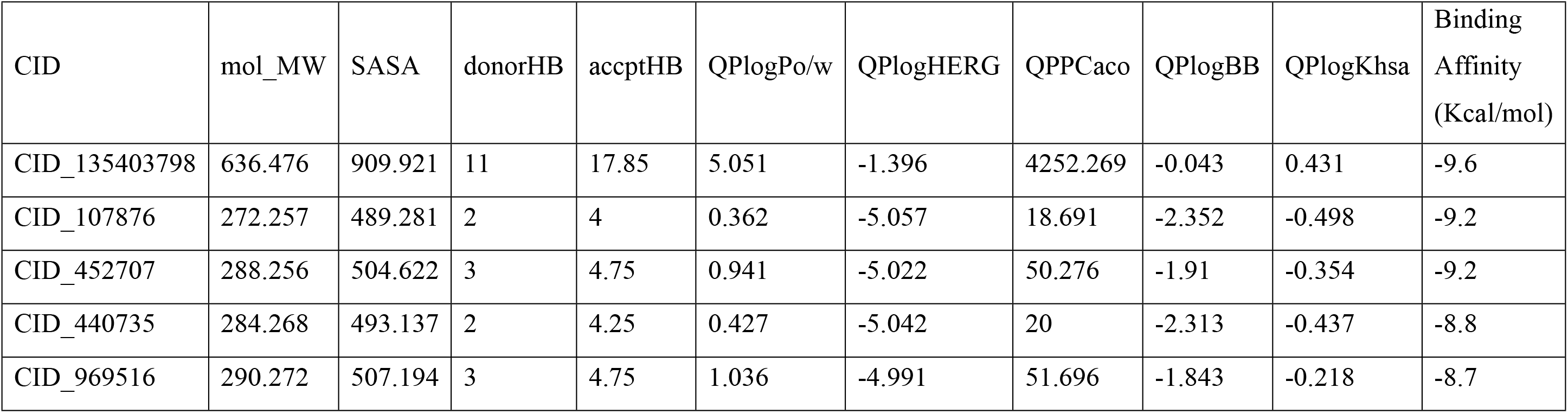
Table showing binding affinity and ADMET analysis of top compounds.

**Figure 2:**
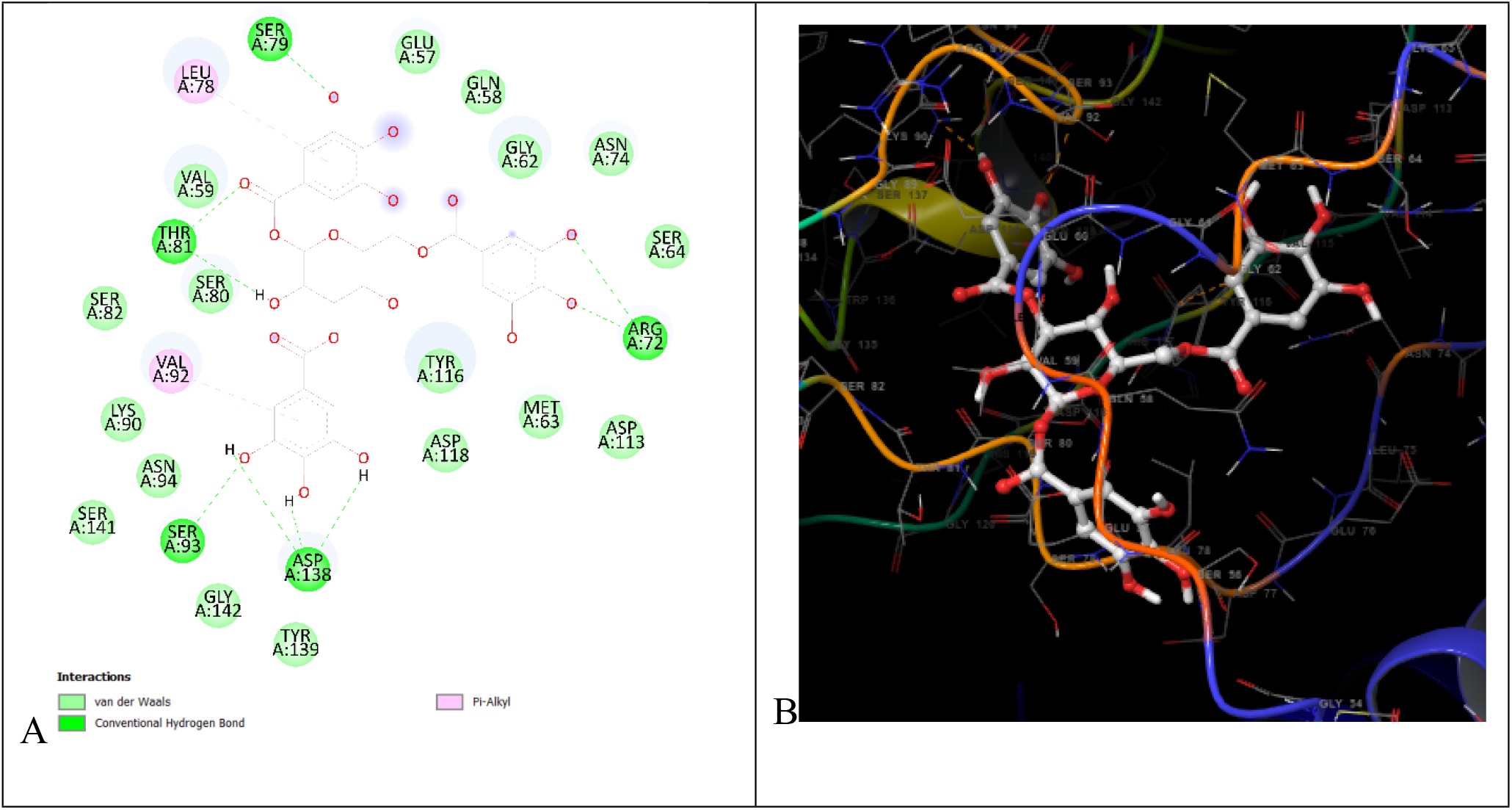
Interactions of lead compound with protein target showing interacting residues of Q997F2-452707 complex in two and three dimensions.

Table 1 displays the molecular weight of the compounds in the “mol_MW” column. The molecular weight range that is considered acceptable is 130.0 to 725.0. The predicted number of hydrogen bonds the solute may establish with water molecules is shown by the variable “donorHB”. The predicted number of hydrogen bonds the solute might accept from water molecules in an aquatic environment is indicated by the symbol “accptHB”. Because this number is an average over several setups, it may not always be an integer. Generally speaking, it should lie between 0.0 and 6.0.

The estimated octanol/water partition coefficient, or “QPlogPo/w” value, indicates how hydrophobic a molecule is. Generally speaking, this number should fall between -2.0 and 6.5. For “QPlogHERG,” the inhibitory concentration (IC50) for HERG K+ channel blockage is shown. When it comes to possible negative consequences, values less than five are deemed worrying. The Caco-2 cell permeability prediction, which gauges how quickly a substance may cross the gut-blood barrier, is represented by the “QPPCaco” number. Values above 500 indicate great permeability, whereas values between 0 and 25 indicate inadequate permeability. “QPlogBB” is a measure of the predicted blood/brain separation ratio that sheds light on a compound’s potential blood-brain barrier crossing capabilities. When using QikProp to anticipate oral medications, the range usually falls between -3.0 and -1.2. This value aids in determining if a substance has the ability to impact the central nervous system. Finally, information on a compound’s binding to human serum albumin may be found using “QPlogKhsa” and “Human serum albumin binding predictions”. The predicted range of these values, which represent the degree of interaction between the chemical and this blood protein, is often between -1.5 and 1.5. Understanding the compound’s pharmacokinetics and its therapeutic uses depends heavily on these factors.

Following the discovery of the lead compound, one particular compound, CID452707 was shown to be the most effective of all the compounds under investigation. The 2D interactions of this high-performing chemical are illustrated in Figure 2, which offers information on its molecular interactions with the target protein. The salient characteristics of this fascinating compound that are critical to its potential as a therapeutic candidate are presented in Table 1. To have a deeper understanding of the interactions and behavior of this optimal chemical combination with the protein target, a 100 nanosecond molecular dynamics simulation was performed. This simulation was performed using Desmond software, and the resulting trajectories were analyzed afterwards. As part of the inquiry, values for root-mean-square-deviation (RMSD) and root-mean-square-fluctuation (RMSF) were calculated. While RMSD provides information on the stability and structural changes of the complex over time, RMSF illustrates the flexibility of the complex’s numerous components during the simulation. A thorough examination of the material’s interactions with the protein provided insight into how the material adheres to and interacts with the target during the simulation. Through the provision of informative information on the chemical’s behavior and potential as a therapeutic candidate, this comprehensive analysis facilitates the evaluation of the compound’s stability, binding qualities, and suitability for further development.

Figure 3 is a graph illustrating the changes in the Root Mean Square Deviation (RMSD) values for the C-alpha atoms in proteins bound to the ligand. RMSD calculates the amount that a protein’s structure deviates from its initial state during a molecular dynamics simulation. Based on the RMSD figure, the complexed protein structure in this simulation reached a stable state at around 10 nanoseconds. This is supported by the RMSD values leveling out and remaining within a range of about 2 Angstroms throughout the duration of the simulation. For predicted proteins such structural stability in proteins is often seen as positive and suggests that the protein maintained its general structure [25]. Still, these changes occurred gradually and with relatively little change. The idea that the protein’s structure was robust and could maintain its integrity throughout the simulation was supported by this outcome. The Ligand Fit to Protein, a measure of how well a ligand molecule fits into a protein’s binding site, did not change during the simulation. The ligand’s RMSD values fluctuated throughout time, suggesting movement or conformational changes, but once equilibrium was attained at five nanoseconds, the ligand’s RMSD showed no appreciable alterations. Ultimately, the RMSD plot demonstrates that the protein-ligand complexes reached a stable state and that, in spite of some oscillations, the ligand and the protein retained their structural integrity during the simulation, demonstrating the stability of the protein structure and suggesting that it is appropriate for additional study.

**Figure 3:**
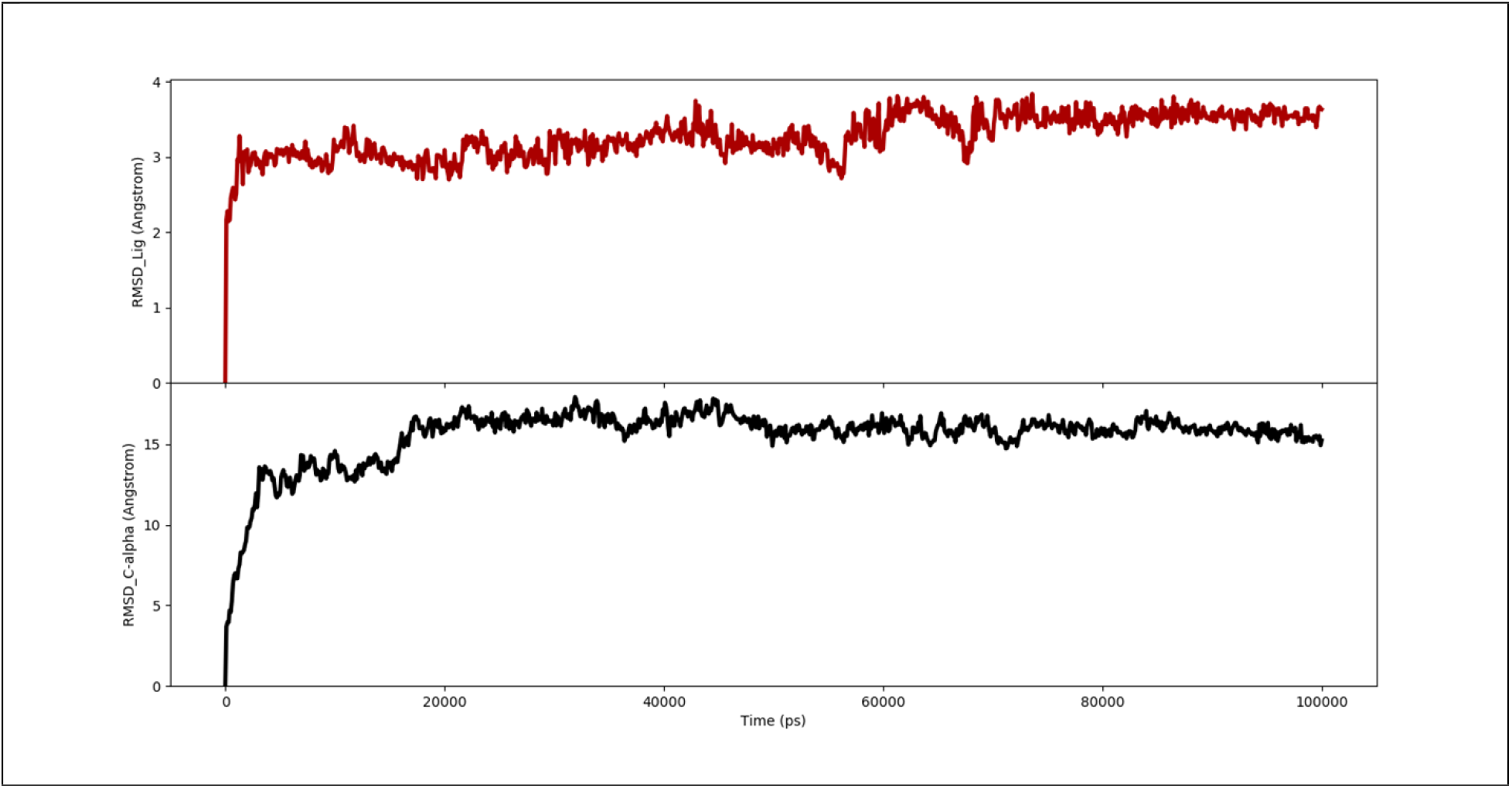
Showing how, over time, the distance between the C-alpha atoms of proteins and the lead molecule has changed. Over the duration of the simulation, the RMSD of the lead molecule is shown in red, whilst the RMSD of the protein target is shown in black.

Figure 4 displays the Root Mean Square Fluctuation (RMSF) values for a protein linked to a ligand. These RMSF values show how much, during a molecular dynamics simulation, certain amino acid residues in the protein structure deviate from their usual locations. The analysis of the molecular dynamics trajectories demonstrates that residues with larger RMSF plot peaks are typically located in flexible regions, such as loops or the protein’s N- and C-terminal ends. This implies that there is more structural variety and dynamic activity in these regions. The stability of the ligand binding to the protein is ascertained by analyzing the RMSF values of the residues in the ligand-binding site. Reduced RMSF values indicate minimal fluctuations in the ligand-protein interaction throughout the simulation for these binding site residues. Figure 5 describes the secondary structural elements (SSE) that were present in the protein during the simulation. It provides a graphic representation of the distribution of beta strands and alpha helices across the protein structure, illustrating the distribution in connection to the residue index.

**Figure 4:**
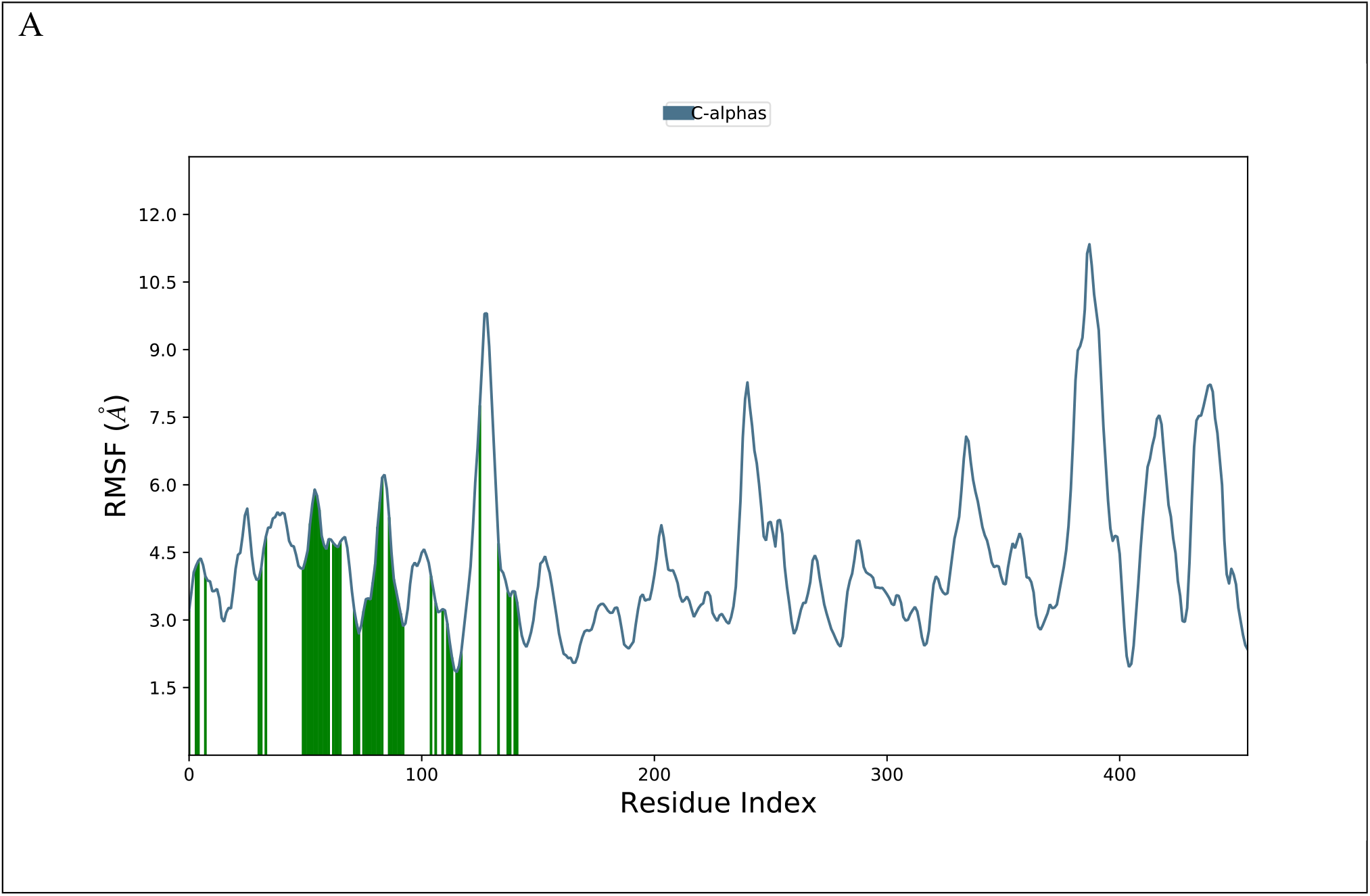
Root Mean Square Fluctuation (RMSF) of a protein in a ligand complex, measured in terms of residues.

**Figure 5:**
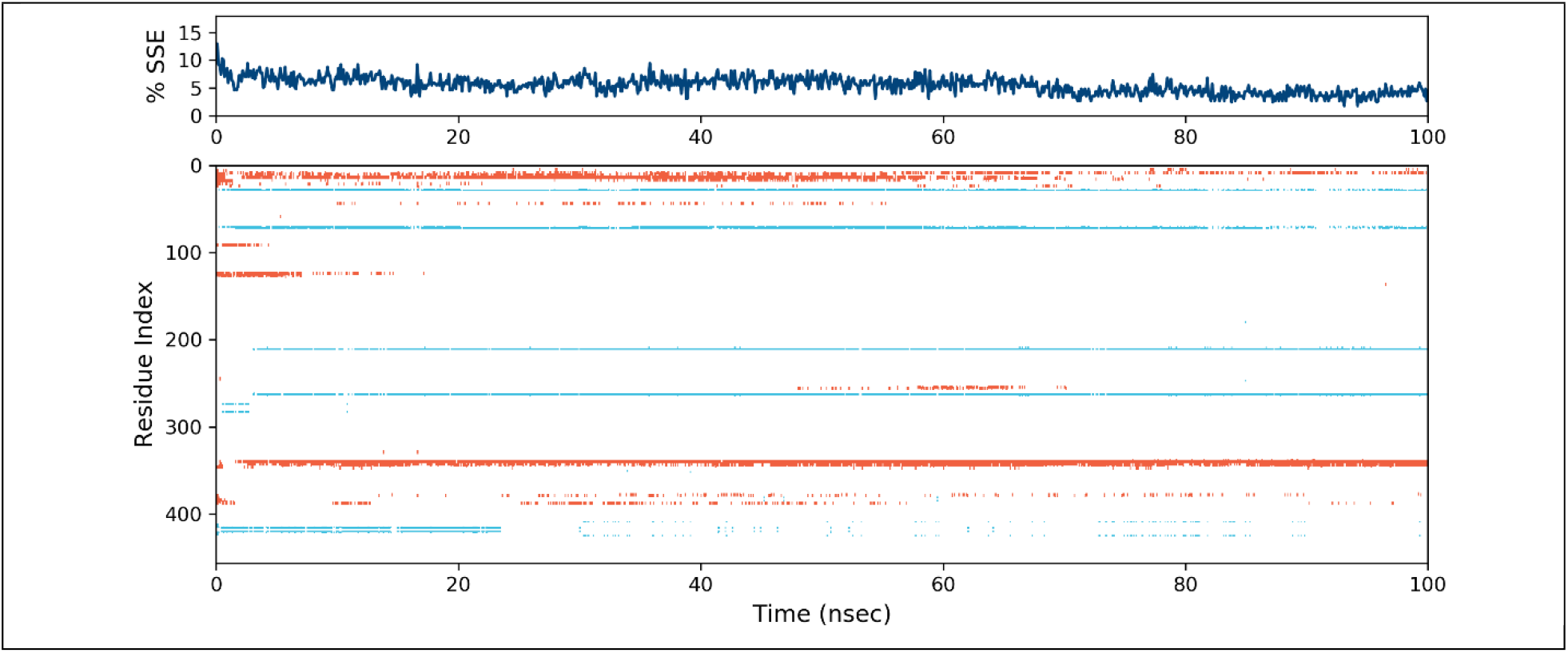
Illustrating the distribution of elements of secondary protein structure within protein-ligand complexes in relation to the position of each residue. Specifically, alpha helices are depicted as red columns, and beta strands are represented by blue columns.

Plotting the SSE against the residue index in the graph illustrates the location of these structural elements inside the protein. Based on the statistical analysis, these constituents account for about 5.49 percent of the secondary structure of complex protein-CID452707. More precisely, alpha-helices account for around 3.34 percent of secondary structural components, whereas beta-strands account for approximately 2.16 percent. Together, these components account for approximately 5.4 percent of the secondary structure of the protein in complex protein-CID452707. Figure 4, which provides a summary of the flexibility of individual residues within the protein, displays more flexible or dynamic regions of the protein based on higher RMSF values. The binding site’s reduced RMSF values indicate that ligand-protein interactions are stable. Figure 5 shows the relative amounts and distribution of beta-strands and alpha-helices inside the protein’s secondary structure. This study provides crucial information on the stability of the protein’s interactions with the ligand and its simulated behavior.

The primary form of interaction that takes place between the protein and the ligand during the molecular dynamics (MD) simulation is represented by hydrogen bonds, as seen in Figure 6. These hydrogen bonds are essential for the ligand’s stability and protein binding. In particular, hydrogen bonding plays a major role in the interactions between certain amino acids and the ligand. Notable amino acids are GLY_54, GLU_57, and ASP_118 in the protein-CID452707complex. These amino acids are especially sensitive to hydrogen bonding interactions and are necessary for the ligand to bind to the protein efficiently. The dynamic changes in the contacts and interactions between ligands and proteins during the experiment are depicted in the timeline in the chart below Figure 6. This timeline allows researchers to monitor the timing and manner of these crucial interactions during the simulation, offering valuable information into the dynamics and stability of the ligand-protein complex. To sum up, Figure 6 illustrates the significance of hydrogen bonds in the interactions that occur between the protein and ligand. It highlights certain amino acids whose role in binding is dependent on hydrogen bonding. The timeline display, which offers a dynamic view on the evolution of these interactions, facilitates a deeper understanding of the complex’s behavior during the MD simulation.

**Figure 6:**
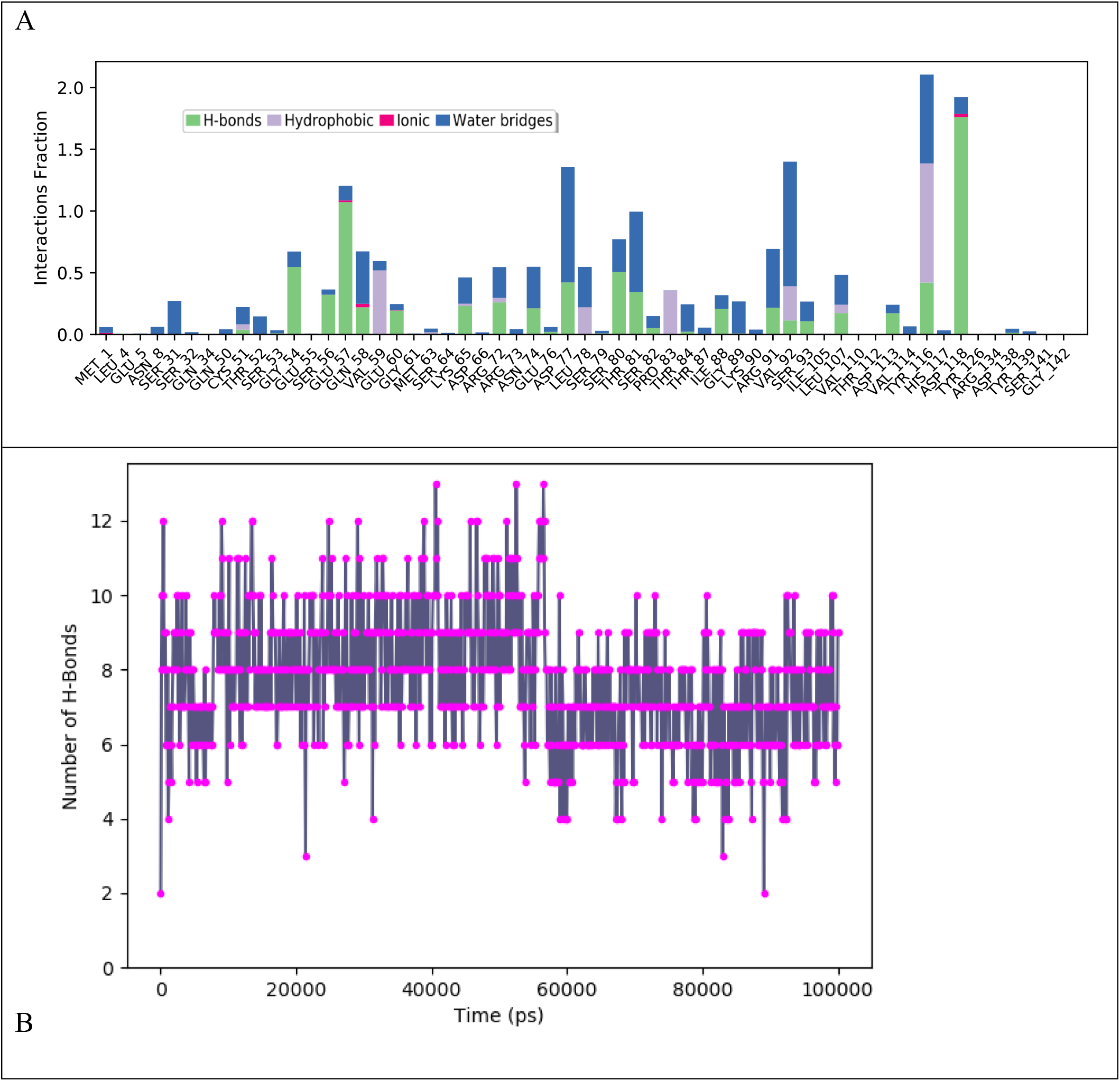
Protein-ligand contact histogram throughout trajectory. A: Histogram of different interaction with proteins in their proportion, B: Number of H-Bonds throughout the time of simulation.

Protein ligand binding affinity was calculated throughout the time of simulation for each frame. Total binding affinity of best ligand with protein is shown in Fig. 7. The average binding free energy between ligand and receptor protein was found to be 47.37 Kcal/mol. That shown robust interations of protein-ligand.

**Figure 7:**
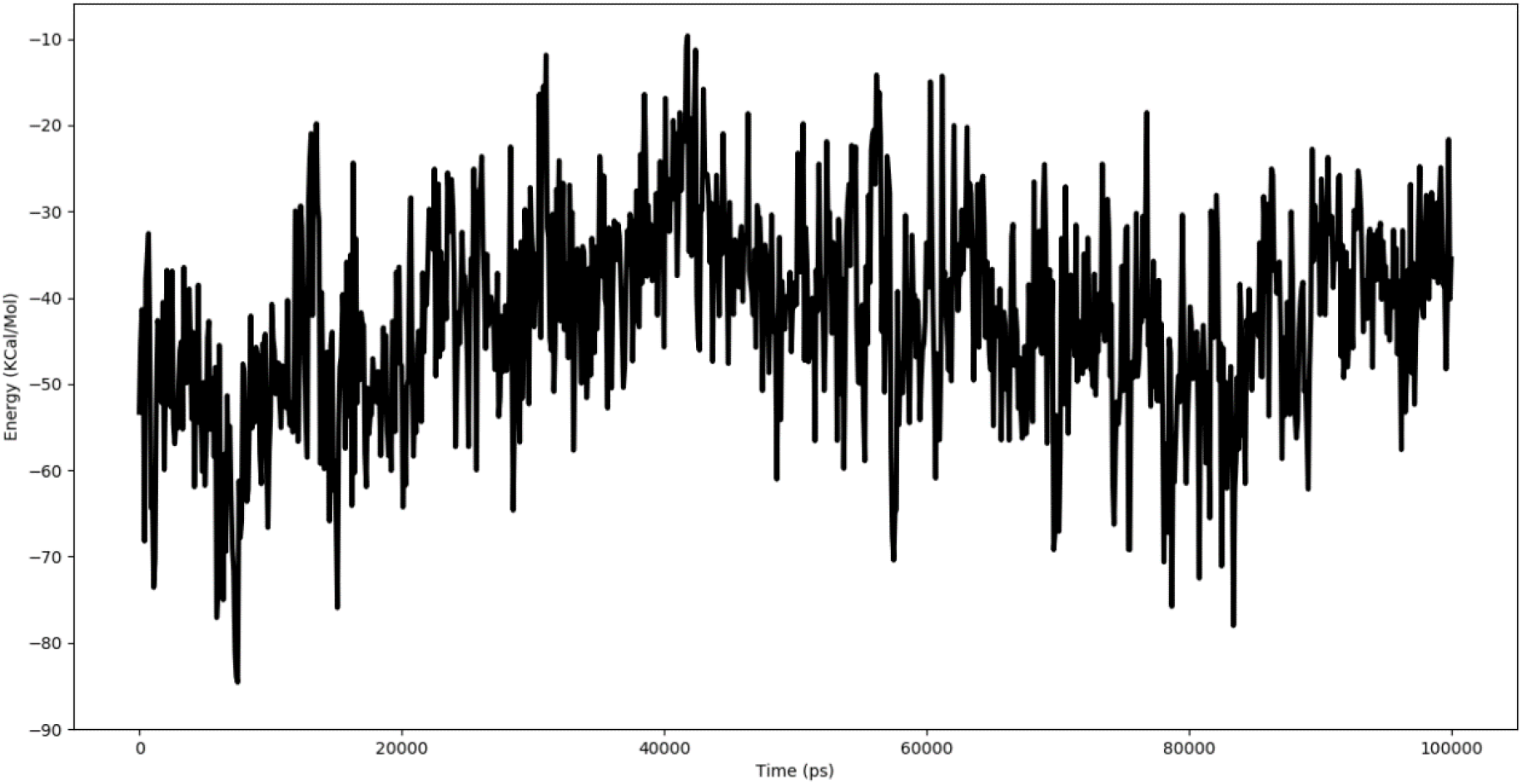
Protein-ligand binding free energy through the time of simulation for each frame.

## Conclusion

Many studies in the field of drug development have been driven by the potential for interdisciplinary methodologies to expedite the process and reduce overall costs. This study’s primary objective was to identify specific target proteins associated with the Nipah virus, which would pave the way for the selection of a lead drug candidate. In order to counteract the lead chemical’s effect on the viral protein, we searched for substances that have this quality. CID452707 possible natural inhibitor, was found. This inhibitor significantly reduces the protein activity at its receptor location. Our rationale is that this material might serve as a platform for developing a medication that selectively targets the Nipah virus without interfering with other biological processes. These findings are extremely promising to scientists and may eventually contribute to the development of a novel treatment for this viral disease.

## Acknowledgment

The authors extend their appreciation to the Deanship of Research and Graduate Studies at King Khalid University, KSA, for this support.

## Data Availability Statement

The data generated during this study are included in this article.

## Ethical Approval

Not applicable

